# Differences in post-mating transcriptional responses between conspecific and heterospecific matings in *Drosophila*

**DOI:** 10.1101/2020.03.25.009068

**Authors:** Yasir H. Ahmed-Braimah, Mariana F. Wolfner, Andrew G. Clark

**Affiliations:** Department of Molecular Biology and Genetics, Cornell University, Ithaca, NY

## Abstract

In many animal species, females undergo physiological and behavioral changes after mating. Some of these changes are driven by male-derived seminal fluid proteins, and are critical for fertilization success. Unfortunately, our understanding of the molecular interplay between female and male reproductive proteins remains superficial. Here we analyze the post-mating response in a *Drosophila* species that has evolved strong gametic incompatibility with its sister species; *D. novamexicana* females produce only ~1% fertilized eggs in crosses with *D. americana* males, compared to ~98% produced in within-species crosses. This incompatibility is likely caused by mismatched male and female reproductive molecules. In this study we use short-read RNA sequencing to examine the evolutionary dynamics of female reproductive genes and the post-mating transcriptome response in crosses within and between species. First, we found that most female reproductive tract genes are slow-evolving compared to the genome average. Second, post-mating responses in con- and heterospecific matings are largely congruent, but heterospecific mating induces expression of additional stress-response genes. Some of those are immunity genes that are activated by the Imd pathway. We also identify several genes in the JAK/STAT signaling pathway that are induced in heterospecific, but not conspecific mating. While this immune response was most pronounced in the female reproductive tract, we also detect it in the female head and ovaries. Our results show that the female’s post-mating transcriptome-level response is determined in part by the genotype of the male, and that divergence in male reproductive genes and/or traits can have immunogenic effects on females.

## 1 Introduction

In internally fertilizing organisms, gamete fusion is often preceded by a complex array of biochemical interactions within the female reproductive tract that mediate reproductive success [1,2]. During copulation, males transfer sperm and a cocktail of seminal fluid proteins (SFPs) that facilitate a variety of post-mating effects that are required for successful fertilization [3–5]. In *Drosophila melanogaster*, several of these SFPs are well-characterized and play important roles in post-mating processes within the female reproductive tract [6–15]. These processes include facilitating sperm storage [16,17], inducing ovulation [18], reducing the female’s propensity to remate [7,19], and affecting female longevity and survival [20,21]. In addition, females undergo dramatic physiological and behavioral changes after mating [22–24]. However, little is known about the female proteins that facilitate these post-mating responses and their role in fertilization success [25–28]. This shortcoming is due in part to the difficulty in isolating and characterizing the interacting female proteins. One approach that is widely used to understand female post-mating reproductive processes is to analyze changes in gene expression after mating [29–39].

The majority of studies on the transcriptional dynamics in females after mating have been conducted in *D. melanogaster*. Overall, these studies show that females typically display a transcriptional response within 3 hours after mating [29], and the peak response occurs around 6 hours after mating [30]. These changes in female gene expression are induced by a combination of transferred SFPs, sperm, and non-ejaculate components of mating [29], the latter potentially including copulatory and behavioral cues. Furthermore, the functional categories of genes that are typically up-regulated in mated females include proteases, protease inhibitors, and immune response genes, which suggests that these classes of proteins play important roles in post-copulatory interactions [29,30,32,39–41,41,42]. Indeed, these functional categories are typically enriched among post-mating response genes in species outside the *Drosophila* genus [33,36–38,43], suggesting broad functional conservation of post-mating processes among insects.

Reproductive interactions between the sexes are often subject to intense postcopulatory sexual selection [44,45]. In species where females can simultaneously store sperm from multiple males, the reproductive tract becomes an arena for intense selective forces that act among ejaculates from different males (*e.g.* sperm competition [46]) and between the sexes (*e.g.* cryptic female choice [47]). Furthermore, reproductive interactions can evolve through coevolutionary arms races, which can result in opposing fitness interests between the sexes (*e.g.* sexual conflict [48]). Taken together, these forces can generate rapid evolutionary changes between diverging lineages and can ultimately lead to reproductive isolation between closely related species [49].

Many closely related species are reproductively isolated at the gametic level (*e.g.* [50–55]), whereby sperm from one species fail to fertilize eggs from the other species. Although the mechanisms that cause these post-mating pre-zygotic barriers are not yet well-understood, they likely involve defects in postcopulatory processes that can be directly impacted by biochemical mismatches between male and female proteins. Thus, species that are reproductively isolated at the gametic level provide a unique opportunity to identify the molecular mechanisms within females that mediate post-copulatory processes. For example, Bono *et al.* [35] used females from the cactophilic species, *D. mojavesis*, to analyze the post-mating transcriptional changes that take place in the female reproductive tract after mating to conspecific (*D. mojavensis*) or heterospecific (*D. arizonae*) males. They found significant perturbations in gene expression in heterospecifically-mated females, indicating failed molecular interactions between male and female proteins that are consistent with the strong gametic incompatibility between this species pair. In addition to being the first of its kind, this study highlighted the utility of using closely related sister species for identifying the post-mating molecular events that are essential for fertilization success.

The *Drosophila virilis* species complex has recently emerged as an ideal system to study the genetic basis of reproductive interactions and their evolutionary consequences [56]. This species group exhibits strong post-mating pre-zygotic reproductive isolation between member species [53,54,56,57], in addition to marked gametic incompatibilities between populations of the same species [54,58–60]. Between the closely related pair within the virilis sub-group, *D. americana* and *D. novamexicana*, fertilization rate is only ~1% in heterospecific crosses when *D. novamexicana* is the female in the cross [54]. This species pair diverged ~0.5 million year ago, and maintains allopatric distributions in the continental United States [61]. Importantly, gametic isolation is the only detectable reproductive barrier between this species pair, suggesting that postcopulatory sexual selection is a particularly strong divergence force in these species.

Here we exploit this system to analyze the post-mating transcript abundance changes in *D. novamexicana* females after mating to *D. novamexicana* (conspecific) or *D. americana* (heterospecific) males. First, we identify candidate female reproductive tract genes based on tissue-biased expression and analyze their functional categories and patterns of molecular evolution. Second, we characterize the transcript abundance landscape in the female reproductive tract, ovaries, and head after con- or heterospecific mating. Finally, we analyze the set of male-derived mRNAs and identify their tissue origin in males. Our results show that female reproductive tract genes with tissue-biased expression are largely slow evolving in this species group, and that the female post-mating response after heterospecific mating is highly distinct from conspecific mating due to consistent up-regulation of stress response genes shortly after mating.

## 2 Methods

### 2.1 Single-pair matings and dissections

The *D. novamexicana* (15010-1031.04) and *D. americana* (SB02.06) strains were maintained at a constant temperature (22^◦^C) in a 12-hr day/night cycle on cornmeal/sucrose/yeast media. Virgin males (*D. americana* and *D. novamexicana*) and females *D.novamexicana* only) were collected under CO_2_ anesthesia within a day after eclosion and housed in single-sex groups of ~20. On day 12 post-ecolsion, individual males and females were paired without anesthesia and mating was observed. *D. novamexicana* females were either mated to a *D. americana* male (conspecific), a *D. americana* male (heterospecific), or were unmated (virgin). All matings were performed in the morning between 8:00 a.m. and 11:00 a.m., and males were removed immediately after mating. Females were subsequently allocated to one of three post-mating dissection time points: 3, 6, and 12 hours post-mating (hereafter 3 hpm, 6 hpm, and 12 hpm). Virgin females were also allocated to the three dissection time points to control for dissection time. At each post-mating time point, conspecific/heterospecific/virgin females were individually anesthetized and immediately dissected in 1xPBS. The lower reproductive tract (bursa, seminal receptacle, spermathecae, and lower oviduct) was extracted, then the ovaries were removed, and finally the head was severed from the thorax; the three tissues were placed in separate tubes containing ice-cold TRIzol. In addition to those three tissues, the gonadectomized carcass was also preserved for the virgin sample (Figure S1). Each virgin and post-mating reproductive tract sample contained three replicates, and ≥100 reproductive tracts were pooled for each replicate of each treatment. The head, ovaries and carcass samples had two replicates and contained pooled tissues from 50 individuals.

### 2.2 RNA isolation, library preparation, sequencing and mapping

Total RNA was extracted from each sample using TRIzol reagent following manufacturer guidelines (Invitrogen, Carlsbad, CA). Single-end, strand-specific mRNA libraries were prepared using the Illumina TruSeq Stranded mRNA library kit (cat. no. 20020594) following manufacturer guidelines. Libraries were sequenced to 100 bp read lengths on an Illumina HiSeq 2500 at the Cornell Biotechnology Resource Center. Raw reads were first processed by filtering clusters from low quality tiles on the flow cell. Subsequently the first 10 bases of each read were clipped, followed by quality-trimming at both ends to a minimum PHRED quality score of 20. Processed reads were mapped to the *D. virilis* transcriptome (FlyBase r1.06) using Bowtie2 v2.2.2 [62]. Finally, a genome-guided *de novo* transcriptome was generated using Trinity r20140717, using reads generated from the virgin and conspecific samples, in combination with male *D. novamexicana* reads from a previous study (SRP100565, [56]).

### 2.3 Differential expression analysis

#### 2.3.1 Tissue-biased genes

To identify genes in *D. novamexicana* with female tissue-biased expression, we used a filtered count matrix (CPM > 5) of male and virgin female tissue samples. The differential expression tests used a design matrix where the focal tissue is compared against all other male and female tissues. Gene-wise differential expression tests were performed by fitting a quasi-likelihood negative binomial generalized log-linear model to the filtered count data and subsequent gene-wise *F*-test, implemented in edgeR (glmQLFit and glmQLFTest, [63]). Female tissue-biased genes are defined as genes that have >2-fold mRNA abundance and FDR <0.01 when compared to other female tissues in pair-wise comparisons. We performed tissue-bias tests using female samples and separately using tissues from the two sexes. We also include an analysis of tissue-biased genes from male reproductive organs that were obtained from a previous study [56].

#### 2.3.2 Post-mating transcript abundance

To analyze the post-mating transcript abundance landscape in females, we analyzed six female reproductive tract samples (3 conspecific and 3 heterospecific), four ovary samples (2 conspecific, 2 heterospecific), and four head samples (2 conspecific, 2 heterospecific). We compared these post-mating transcript abundance states in each tissue to their respective unmated control samples, in addition to comparisons between the conspecific and heterospecific post-mating samples. For each tissue type, we created a subsetted matrix that only included the tissue to be analyzed and filtered the genes to only include rows where the CPM value was >5 in at least 3 (female RT) or 2 (head and ovary) replicate samples. For the differential expression analysis we implemented removal of erroneous variation between samples using the RUVseq package [64] with *k* = 2 (Figure S3A). We included the residuals from this model in the design matrix that was used to fit a quasi-likelihood negative binomial generalized log-linear model to the count data (edgeR). We then performed differential expression (DE) contrasts between the con- and heterospecific post-mating samples at each time point or between post-mating samples and the virgin sample and classified significantly differentially expressed genes as those that were up- or down-regulated by >2-fold and FDR<0.05.

### 2.4 Transcript annotations and gene ontology (GO) analysis

Custom annotations of the *D. virilis* genome (FlyBase version 1.06) were produced in a previous study [56]. These annotations included UniProt orthologies and orthologs against the *D. melanogaster* genome that were checked against the FlyBase orthology calls. These annotations also included gene ontology (GO) associations for protein sequences that contain an orthologous hit in the UniProt database. GO ontology enrichment analyses were performed using the GOseq R package [65].

### 2.5 Population genetic analysis

The rate of nucleotide and amino acid substitution between species in the *D. virlis* group was calculated using custom Perl scripts and BioPerl libraries (https://github.com/YazBraimah/cbsubin/blob/master/SAPA.pl). Whole-genome DNA-seq data from the four group members (*D. virilis, D. lummei, D. americana* and *D. novamexicana*) was aligned to the *D. virilis* genome and known coding gene sequences (CDS) were extracted. We then generated multiple sequence alignment for each CDS and performed pairwise dN/dS between *D. americana* and *D. novamexicana*, and calulcated *ω* (a close proxy for *dN/dS* [66]) across the subgroup phylogeny using PAML [66]. We also used the multiple sequence alignments to calculate the pair-wise percent conservation of amino acids from cDNA sequences using our custom Perl script. Finally, we performed the “branch-site” test along each branch of the phylogeny to test for the impact of natural selection on each gene [67]. To identify genes with a significant signature of positive selection, we performed a likelihood ratio test (LRT) between a model with *ω* = 1 and a model where *ω* is estimated from the data, and derived *p* values using the *χ*^2^ distribution.

### 2.6 Analysis of transferred paternal mRNAs

To identify paternally transferred mRNAs we first identified annotated *D. virilis* genes in our dataset that show higher abundance at the earliest post-mating time point (3 hpm) in either the conspecific or heterospecific crosses, and only analyzed genes that do not show a decline in abundance at subsequent time points. We then separately identified the *de novo*-assembled *D. novamexicana* transcripts that show higher abundance at 3 hpm and subsequent gradual decline. We used BLAST results between those two sets to identify the overlap between them and used the overlap set for downstream analysis. We then analyzed single nucleotide polymorphisms (SNPs) that are present/absent in the *de novo*-assembled transcripts and identified whether mapped reads from the heterospecific 3 hpm sample contained distinct SNPs; if there is a mismatch between reads from the heterospecific sample and the assembled *de novo* transcriptome, we infer that these reads—and thus, the mRNA that produced them—are paternally derived.

### 2.7 Data and script availability

The Illumina sequence reads are available through the Sequence Read Archive (SRA) under project accession PRJNA611072. The processed data files and analysis scripts are available through a GitHub repository: github.com/YazBraimah/DnovPmRNAseq.

## 3 Results

### 3.1 Evolutionary dynamics of female reproductive tract genes

The female reproductive tract is the site of complex interactions between male ejaculate proteins/sperm and female reproductive tract components. It is thus a hotbed of mutually beneficial—or conflicting—evolutionary dynamics that can drive species differentiation or co-evolution between male and female reproductive genes. It is widely appreciated that male reproductive genes tend to diverge rapidly between species [68,69], but the evolutionary dynamics of female reproductive genes are less clearly understood, owing in part to the difficulty in defining the set of female reproductive proteins that specifically interact with male proteins. In our first analysis we used the unmated female tissue samples to identify female reproductive tract-biased genes to characterize their functional attributes and evolutionary dynamics.

#### 3.1.1 Female reproductive tract genes are enriched for proteolytic enzymes and membrane-bound receptors

To analyze female reproductive tract (fRT) genes in *D. novamexicana*, we defined such genes based on expression bias in the lower female reproductive tract (bursa, spermathecae, seminal receptacle, and oviduct) compared to other female and male tissues. Specifically, we defined fRT genes as those that show >2-fold mRNA abundance (FDR≤0.01) relative to ovaries, head, gonadectomized female carcass, and male tissues. This classification yielded 148 fRT-biased genes (Figure 1A). We performed gene ontology (GO) enrichment analysis on this set of genes and found that they are enriched for serine-type peptidases, suggesting that the fRT plays roles in dictating proteolytic cleavage of male and/or female compounds [70,71] (Figure 1B). All ten proteolytic enzymes are almost exclusively expressed in the fRT, and contain six trypsins, 3 serine proteinases (*Stubble*-like) and one collagenase (Table S1). One of the trypsins is orthologous to the *D. melanogaster Send1/Send2* proteins, which are specifically expressed in the secretory cells of the spermatheca [27] (Table S1). We also found a significant enrichment of plasma membrane-bound proteins, some of which are known receptors that might bind male-derived compounds. For example, one of those receptor proteins is the *D. melanogaster* ortholog of a gustatory receptor (*Gr39a*), which shows fRT-specific expression and has likely been co-opted into a reproductive function in this species (Table S1).

**Figure 1.**
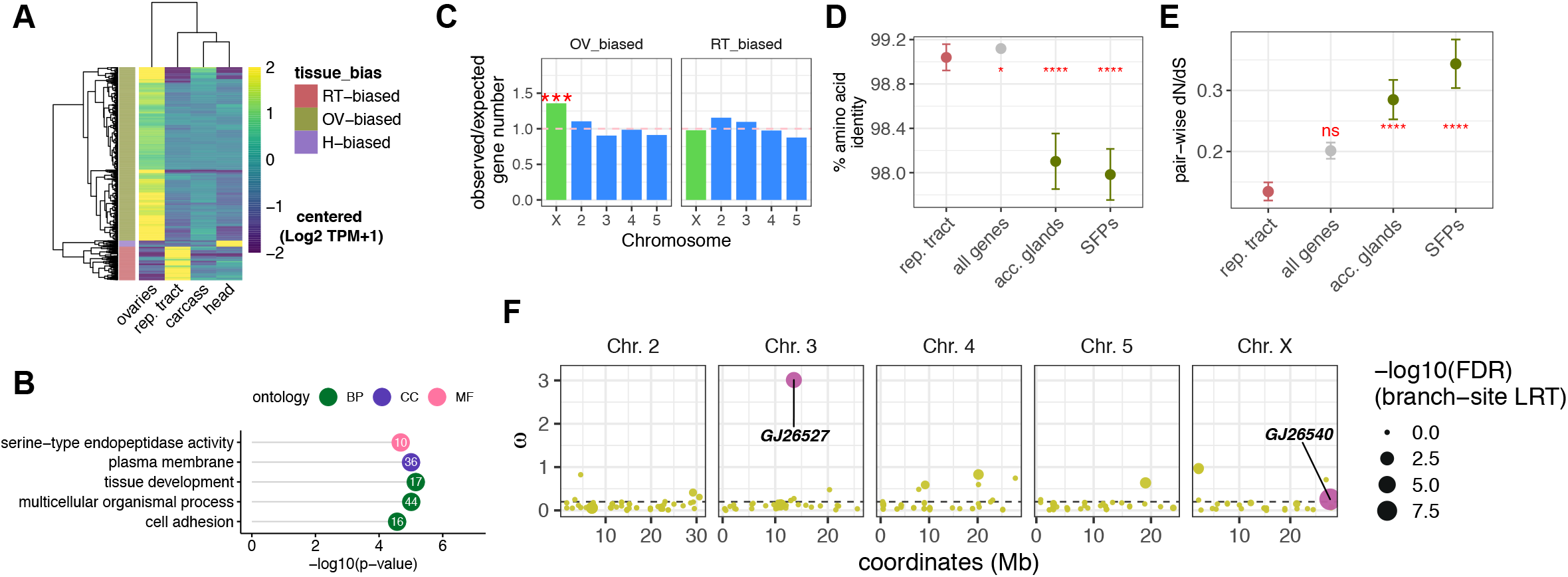
Functional and evolutionary dynamics of female reproductive tract genes. **(A)** Heatmap of female tissue-biased genes, with tissue categories shown on the left and key on the right (RT: lower reproductive tract; H: head; OV: ovaries). **(B)** Lollipop plot of gene ontology enrichment of fRT-biased genes. The number of fRT genes within each enriched category is indicated within the circles. (BP: Biological Process, CC: Cellular Component, MF: Molecular Function). **(C)** Chromosomal distribution of ovary-biased and fRT-biased genes. The horizontal dashed-line at *y* = 1 indicates random expectation of observed/expected number of genes on a given chromosome. **(D)** Amino acid conservation between *D. americana* and *D. novamexicana* reproductive genes. “SFPs” are the subset of accessory gland-biased genes that contain a predicted signal sequence. Error bars represent standard error. **(E)** Average pair-wise non-synonymous to synonymous substitution rate (dN/dS). **(F)** Gene-wise *ω* (a proxy for dN/dS) for fRT genes along the five major chromosomes. The dotted line indicates the genome average *ω* value (0.2). The size of each dot indicates the −log10(FDR) derived from the LRT under a *χ*^2^ distribution. The two significant genes (FDR<0.05) are indicated. (*: *p* ≤ 0.05, **: *p* < 0.01, ***: *p* < 0.001, ****: *p* < 0.0001)

The chromosomal distribution of reproductive genes can often be informative with regards to the evolutionary forces that impact them. For instance, male reproductive proteins—particularly seminal fluid proteins—tend to be under-represented on the X chromosome [4,56]. We found that fRT-biased genes do not show a biased distribution on the X chromosome, suggesting that their chromosomal distribution is not affected by sex-specific selection (Figure 1C). However, ovary-biased genes are over-represented on the X chromosome (Figure 1C; [72]).

We were surprised to find that, if we do not include male reproductive tissues in the expression bias analysis, several fRT-biased genes also show reproductive tissue-biased expression in males (Figure S2A). To explore this further, we analyzed fRT-biased genes using female tissues only, and examined the overlap with male reproductive tissue-bias (accessory glands, ejaculatory bulb, and testes). Strikingly, we found that 14, 31, and 20 fRT biased-genes are also biased in the accessory-glands, ejaculatory bulb, and testes, respectively (Figure S2B). Several of these genes show exclusive expression in the fRT in females and a corresponding male reproductive tissue (examples in Figure S2C).

These results show that our classification of female reproductive tract genes using mRNA expression bias can provide robust candidates for analyzing female reproductive genes. These results also show that using sex-bias as a criterion for defining reproductive genes might miss a set of reproductive genes that are shared between the sexes.

#### 3.1.2 Female reproductive tract genes show a reduced rate of sequence divergence compared to male accessory gland-derived genes

One of the hallmarks of reproductive genes is their rapid divergence between species [68], yet the extent to which this phenomenon holds for female reproductive genes remains understudied. Here we analyzed the rate of nucleotide divergence by examining (1) conservation of amino acid sequence between *D. americana* and *D. novamexicana*, (2) the average pair-wise ratio of non-synonymous to synonymous substitutions between *D. americana* and *D. novamexicana*, and (3) evidence of positive natural selection using the “branch-site” test implemented in PAML [67].

First, we found that fRT genes are highly conserved but are slightly less conserved than the genome average (Figure 1D, *p* = 0.024; Mann-Whitney U test). In contrast, male accessory gland genes, especially those that contain a predicted secretion signal, are less conserved compared to fRT genes and the genome average (Figure 1D, *p* ≪ 0.0001; Mann-Whitney U test). Second, the average ratio of non-synonymous to synonymous substitutions is lower in fRT genes compared to the genome average, although this difference is not significant (Figure 1E, *p* = 0.2; Mann-Whitney U test). It is, however, significantly lower than male accessory gland genes and predicted SFPs (Figure 1E, *p* ≪ 0.0001; Mann-Whitney U test). Finally, we performed the “branch-site” test in PAML to examine evidence of positive selection at any of the 148 fRT-biased genes. We performed the test using the core *D. virilis* species sub-group (*D. americana*, *D. lummei*, *D. novamexicana*, and *D. virilis*), and found that only two genes show evidence of a significant signature of positive selection. One gene (*GJ26540* ; LRT = 52.3, FDR = 6.2×10^−10^) resides on the X chromosome, has a substitution ratio that is equal to the genome average, and has no known ortholog or function. The other gene (*GJ26527* ; LRT = 26.1, FDR = 0.0001) resides on chromosome 3, has the highest *ω* ratio among fRT-biased genes, and is orthologous to Titin, which is a muscle-associated protein. Thus, we conclude that fRT genes tend to be conserved relative to male reproductive genes in this species group, but a small subset of genes could experience rapid bouts of selection.

### 3.2 Post-mating transcript abundance changes in females after con- or heterospecific matings

Females undergo a variety of physiological changes after mating, and these changes are induced in part by male-derived molecules that are transferred during copulation [13,24]. Many of these physiological changes are likely driven by changes in gene expression that trigger cascades of concomitant biochemical changes throughout female tissues. Unfortunately we still do not fully understand how the molecular interplay between male ejaculate components and the fRT results in these post-mating changes, and how these changes affect fertilization success. To ameliorate this problem, here we sought to analyze the post-mating transcript abundance changes that occur in *D. novamexicana* females after mating with a conspecific male or a heterospecific male. This allows us to identify the impact of divergent male molecules (SFPs, other ejaculate molecules, and sperm proteins) on post-mating transcript abundance in females, and may shed light on the importance of these changes to the observed fertilization incompatibility in heterospecific crosses between *D. novamexicana* and *D. americana*.

#### 3.2.1 Heterospecifically-mated females show a distinct transcript abundance profile in the fRT compared to conspecifically-mated females

We collected mRNA abundance data from the fRT after two distinct mating conditions (conspecific or heterospecific), at three different time points after mating (3 hr, 6 hr, and 12 hr), and from unmated females (virgin) as a control. First, after filtering transcripts with low counts, we examined the grouping of replicates within each sample using a Pearson correlation matrix to assess the suitability of our downstream analyses, and found that replicates do indeed cluster as expected (Figure S3B). Next, we examined replicate and sample groupings by constructing a multidimensional scaling (MDS) plot of fRT data using the filtered counts, and observe distinct clustering of post-mating samples based on cross type and post-mating time point (Figure 2A). Specifically, early post-mating time points (3 hpm and 6 hpm) were clearly distinct from the virgin control sample, and showed the strongest difference between the two cross types. At 12 hpm the two cross types were somewhat congruent and clustered closer to each other than the two preceding time points. These observations suggest that, shortly after mating, the transcript abundance profile between females mated to a conspecific or a heterospecific male are highly distinct.

**Figure 2.**
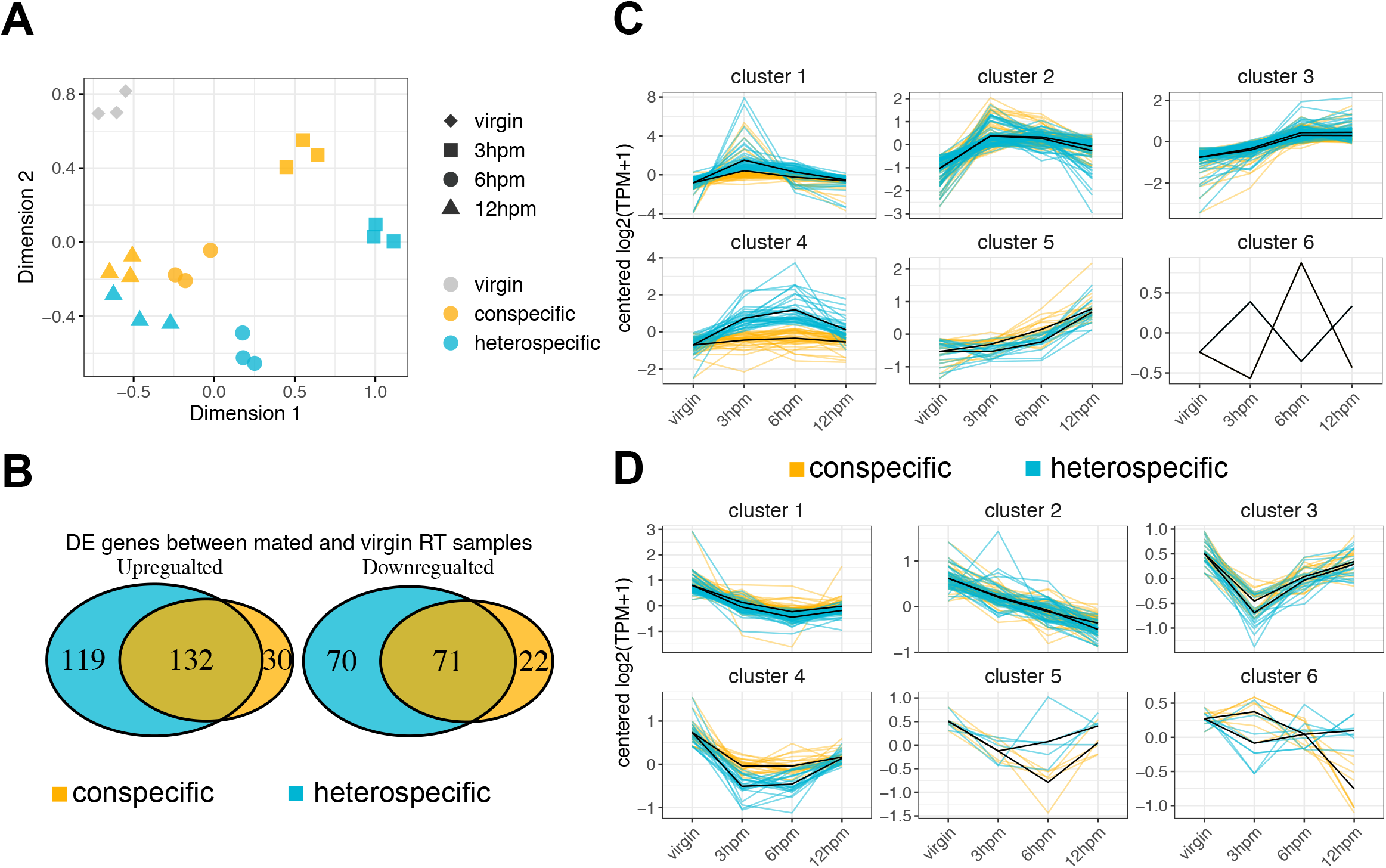
Post-mating transcript abundance profiles differ between conspecific and heterospecific crosses. **(A)** Multi-dimensional scaling (MDS) plot of unmated and post-mating fRT samples, using the first two dimensions. The key on the right indicates the post-mating time-point or unmated sample (shape) and the cross type (color). **(B)** Venn diagram depicting the overlap between all post-mating DE genes. **(C)** and **(D)** *k*-means clustering of post-mating DE genes. Conspecific and heterospecific transcript abundance responses are plotted together and are distinguished by color (see key). Solid black lines represent the average abundance profile for each of the two cross types.

Next, we performed pair-wise differential abundance analysis between each post-mating time point and the virgin control sample for both cross types separately. We define significantly differentially expressed (DE) genes as those with >2-fold change in abundance (FDR < 0.05) in the post-mating sample relative to the virgin control. We found many more genes that are up-regulated after mating (281) than down-regulated (163) (Figure 2B). Furthermore, the number of DE genes in the heterospecific mating samples is greater than the conspecific samples: 251 up-regulated and 141 down-regulated in heterospecific matings, 162 up-regulated and 93 down-regulated in conspecific matings, and 132 shared up-regulated and 72 shared down-regulated in both cross types (Figure 2B).

We performed GO analysis for all up-regulated genes and found several enriched GO categories (Figure S4A; FDR < 0.05). Prominent among these were terms associated with the proteasome complex, which is involved in the degradation of poly-ubiquinated proteins in the nucleus and the cytosol [73]. Seventeen genes that code for various subunits of the proteasome core complex gradually increase in abundance after mating to either a con- or heterospecific male, however the magnitude of increase is markedly higher among heterospecifically-mated females than conspecifically-mated ones (Figure S4B). Other enriched GO terms represent other proteolytic processes and the immune response (see below). Only two GO terms are significant among down-regulated genes: oxidoreductase and alanine-glyoxylate transaminase activity.

To further explore the post-mating expression profiles across the three time-points, we performed separate *k*-means clustering of the up- and down-regulated DE genes, with *k* = 6 (Figures 2C and 2D, respectively). This analysis revealed that the expression profiles between the con- and heterospecific cross types are largely congruent, but two clusters in the up-regulated set (cluster 1 and cluster 4) contained several genes that show distinct expression profiles between the con- and heterospecific cross types (Figure 2C). We performed GO enrichment analysis on the up-regulated clusters and found that clusters 2, 3, and 4 were enriched for several GO categories. Cluster 2 contained several terms related to stress response mechanisms (*e.g.* response to hypoxia, topologically incorrect proteins, and acid chemicals), which are primarily driven by several heat shock proteins that are up-regulated at 3 hpm in both con- and heterospecific crosses (Table S3). Cluster 3 was enriched for terms related to proteolytic activity and the proteasome complex, which reflect the overall composition of all up-regulated transcripts. Notably, cluster 4 primarily contained an enrichment of immune response genes, and these appear to show the most distinct abundance response between con- and heterospecific cross types.

To probe the abundance differences between the con- and heterospecific cross types directly, we performed DE analysis between the two post-mating samples at each time-point. We identified 65 genes that show significantly higher abundance in the heterospecific samples across all time-points, and only 15 that show significantly higher abundance in the conspecific sample (Figure S4C). Of the 65 genes that show higher abundance in the heterospecific cross, 47 are significant at the earliest time-point (3 hpm). We performed GO enrichment analysis on these 65 genes and recovered significant enrichment of immune response terms (Figure S4D). These immune genes include the NF-*κ*B signaling protein, Relish, which is a transcription factor that is a master regulator of immunity through the Imd pathway [74] (Figure 3A). Several immune effectors that act as antimicrobial peptides (Attacin, Cecropin, Defensin, and Diptericin) are also up-regulated distinctly in the heterospecific cross, and show little or no response after conspecific mating (Figure 3A). Furthermore, other classes of immune-related genes show a distinct response in the heterospecific cross, including a recognition protein with beta-glucan binding activity (Gram-negative bacteria binding protein, GNBP) and a coagulation effector protein (Tiggrin) (Table S3).

**Figure 3.**
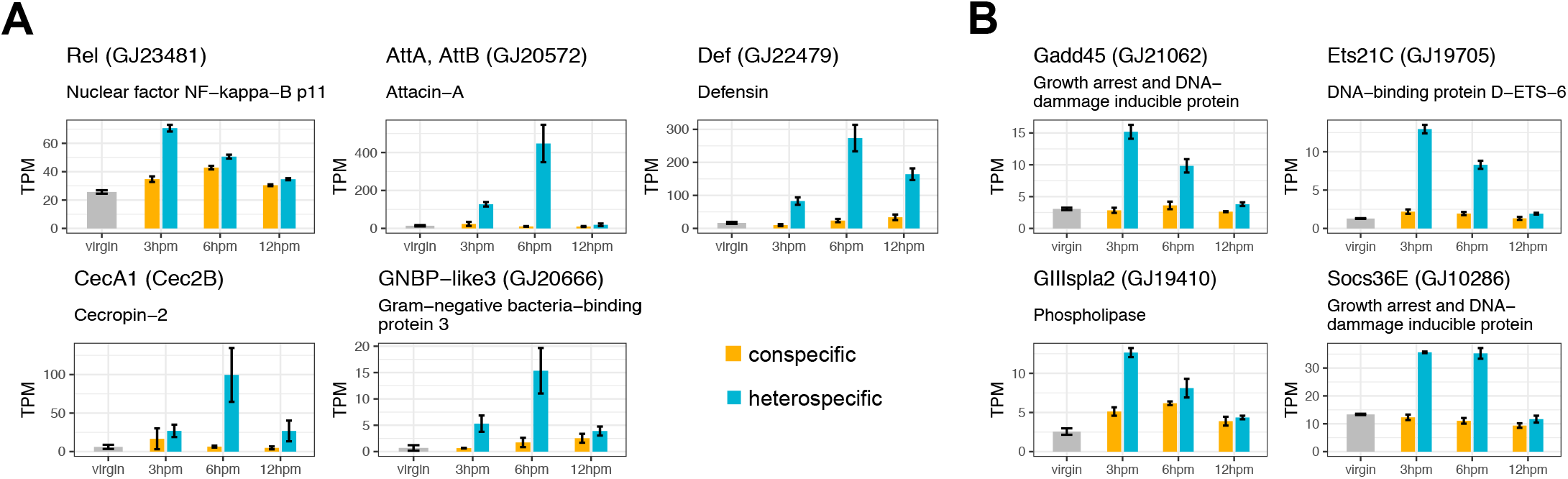
Misregulated post-mating genes in the fRT. A subset of **(A)** immune response genes and **(B)** stress response genes that are up-regulated in the heterospecific cross but remain largely unchanged in the conspecific cross. The *D. melanogaster* ortholog gene name is shown, followed by the *D. virilis* gene name and a gene description derived from a SwissProt database search. Error bars represent standrd error.

We also found that several genes that are involved in some stress response pathways in *Drosophila* are up-regulated in the heterospecific cross but mRNA levels remain largely unchanged in the conspecific cross. These include a negative regulator of the Janus Kinase/Signal Transduction and Activator of Transcription (JAK/STAT) signaling pathway, *Socs36E*, which shows a ~3-fold increase in abundance at 3 hpm and 6 hpm only after heterospecific mating (Figure 3B). Another stress response gene is a phospholipase, *GIIIspla2*, which acts as a downstream target of various stress response pathways and metabolizes phospholipids.

These results show that the two cross types largely induce congruent changes in transcript abundance that are likely required for normal post-mating processing, but that the heterospecific cross induces additional abnormal changes in transcript abundance that indicate a heightened stress response.

#### 3.2.2 Several genes are mis-regulated in ovaries after heterospecific mating

We analyzed transcript abundance changes in ovaries and female heads after con- or heterospecific mating at 6 hpm. In *D. melanogaster*, mating increases oogenesis and stimulates ovulation in part through the action of transferred male SFPs (*e.g.* Sex Peptide and *ovulin*; [14,75,76]). We therefore reasoned that heterospecific mating could cause misregulation in ovaries.

Our analysis of the ovaries revealed that, indeed, transcript abundance profiles of mated female ovaries at 6 hpm are distinct between the two cross types: conspecifically-mated female samples cluster separately from heterospecifically-mated females (Figure S5A-B). Differential expression tests revealed that more genes are differentially abundant after conspecific mating relative to virgin (96 up-regulated and 25 down-regulated; Table S4), compared to heterospecific mating relative to virgin (23 up-regulated and 4 down-regulated; Table S4). However, when we compared the two post-mating samples against each other (conspecific vs. heterospecific) we found that only 11 genes show higher abundance in the conspecific sample. Of the 11 genes with higher abundance in the conspecific sample, three are orthologous to Cytochrome C oxidase subunit 1 or 2 (Figure 4). In addition, one of the 11 genes is orthologous to the *D. melanogaster* actin gene, *Act79B*, which localizes to the actin cytoskeleton and is often expressed in muscle tissue [77]. Conversely, only one gene—an fRt-biased, textilinin-like protease inhibitor—shows higher abundance in the heterospecific sample compared to the conspecific sample. The *D. melanogaster* ortholog of this gene has not been characterized, but is a predicted kunitz-type serine protease inhibitor that is predicted to localize to the extracellular matrix (FlyBase.org).

**Figure 4.**
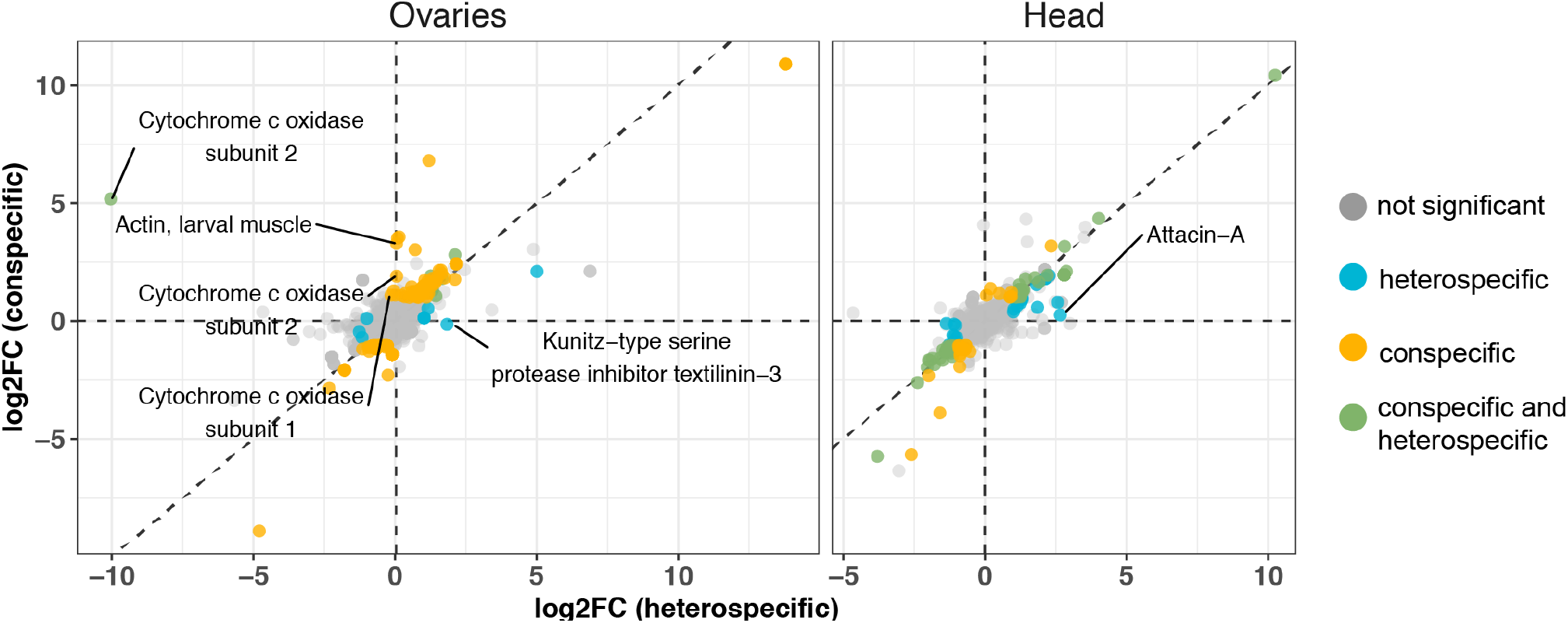
Log-fold change in the ovaries and female head at 6 hpm. The log2 fold-change of heterospecific crosses is represented on the *x*-axis, and the log2 fold-change of conspecific crosses is represented on the *y*-axis. Each point represents a single gene, and the DE status is indicated by color (see legend). “Normal” gene data points are defined as those that are DE in both the conspecific and heterospecific cross when compared to the unmated control, and show a log2 fold-change in the same direction. “Conspecific” and “heterospecific” gene data points refer to those that are significantly DE between the conspecific post-mating sample—and not the heterospecific sample— and the unmated control, or between the heterospecific sample—and not the conspecific sample—and the unmated control, respectively. Genes that are significantly differentially expressed between the conspecific and the heterospecific sample are indicated by text (color represents the cross type in which the gene is higher in abundance).

We performed GO enrichment analysis of up-regulated genes in the ovaries and, as expected, recovered significant terms associated with developmental processes involved with reproduction and vitelline membrane formation (Table S5). We also recovered terms associated with the immune response, and this enrichment was driven by ten genes—*e.g.*, two peptidoglycan-recognition proteins, thioester-containing protein, and Cathepsin—that were similarly up-regulated in the con- and heterospecific cross. Interestingly, one antimicrobial peptide, Cecropin 2B, is up-regulated in the conspecific sample after mating (∼2k-fold) but massively up-regulated after the heterospecific cross (~17k-fold; Table S4).

These findings suggest that heterospecific mating induces distinct post-mating transcriptional changes in ovaries.

#### 3.2.3 A single antimicrobial peptide is detected as mis-regulated in the female head after heterospecific mating

Insect females undergo various behavioral changes after mating that might reflect underlying transcriptional changes in the head. Indeed, female *D. novamexicana* up-regulate 51 genes and down-regulate 56 genes at 6 hr post-mating in head tissue (Table S6). Notably, the post-mating transcript abundance changes in the female head are almost identical after con- or heterospecific mating and replicates do not cluster by cross (Figure S5B). We sought to examine whether the two cross types induce different transcript abundance patterns at 6 hpm, but found that only one gene shows a significant increase in the heterospecific cross but remains unchanged in the conspecific cross: the antimicrobial peptide Attacin-A. This gene is also up-regulated in the fRT after heterospecific mating but not conspecific, suggesting that the mechanism of induction could be organism-wide.

### 3.3 Males transfer testes-expressed mRNAs to females during copulation

Two studies that examined post-mating gene expression in *D. mojavensis* and *Aedes aegypti* females have shown that some mRNAs that may appear to be up-reglated in the fRT after mating are derived from male reproductive tissues [35,37]. In both cases these male-derived mRNAs were highly expressed in the male accessory glands, and some have been characterized as SFPs [37]. Although the functional significance of these male-derived mRNAs in the fRT is not clear, it may indicate some as yet unknown male contributions to the female through the ejaculate.

We identified several genes that were “up-regulated” after mating and that exhibited high expression in male reproductive tissues. This prompted us to examine whether these transcripts are transferred from the male during copulation. First we identified all mRNAs that were up-regulated at 3 hpm and gradually decreased in abundance in subsequent time points, which is an abundance pattern that indicates transfer from males and subsequent decay or usage (see cluster 1 and 2 in Figure 2C). We confirmed—using species-specific single nucleotide polymorphisms (SNPs)—that 13 of these mRNAs are male-derived (Figure 5, Table S6). Notably, three of these encode heat shock proteins (*Hsp23*, *Hsp68*, and *Hsp70Ab*) that show testis-biased expression but have relatively low normalized abundance (transcripts per million, or “TPM” ~7-60, Table S4). All but two of the remaining genes also show strong testis-biased expression but high relative normalized abundance (TPM ~280-61,000, Table S6).

**Figure 5.**
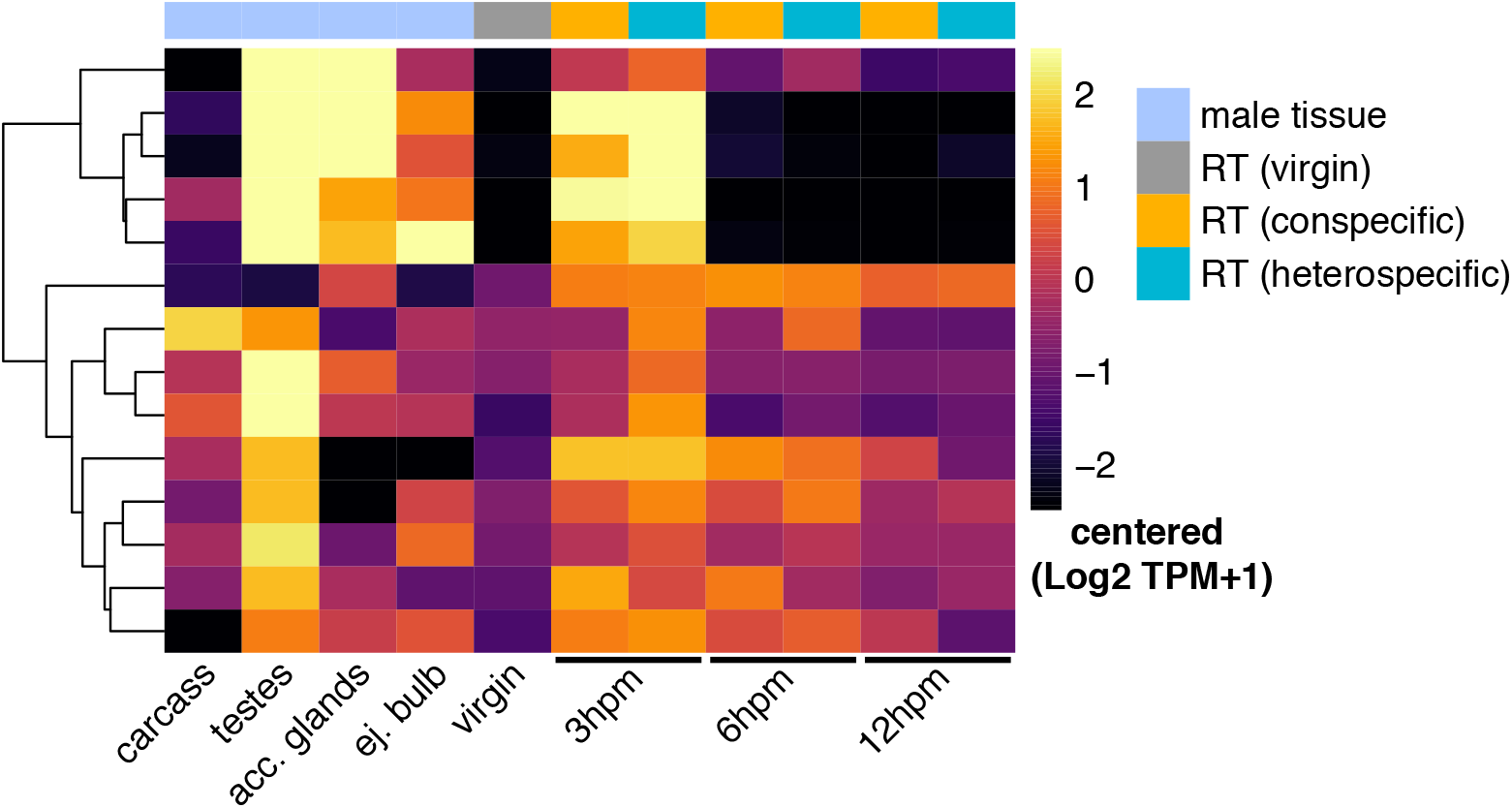
Transcript abundance heatmap of male-derived mRNAs. Sample origin is indicated below each column and the tissue category is indicated by the colored annotation bar on top (see legend).

These results show that paternally-derived mRNA transcripts that are found in the mated fRT can originate from testes, which shows that testes can contribute non-sperm components to the ejaculate.

## 4 Discussion

Here we investigated the mRNA abundance patterns within the reproductive tract of unmated *D. novamexicana* females and females mated to either conspecific or heterospecific (*D. americana*) males. This species pair is well-suited for genetic investigations of postcopulatory interactions as they have diverged recently (~0.5mya) and have evolved strong gametic incompatibilities [54,61]. These incompatibilities manifest as reduced fertilization and premature loss of sperm from storage after heterospecific mating. Because these incompatibilities are likely caused by a mismatch between the male ejaculate and the female reproductive tract, we sought to identify incongruent transcript abundance changes in females after mating with conspecific or heterospecific males to identify potential molecular mechanisms within the female that mediate reproductive success. We also used the mRNA abundance data to identify fRT-biased genes, as these are likely to have specialized reproductive functions.

One of the most pervasive patterns in molecular evolution is the rapid divergence of reproductive genes in a wide variety of organisms [68,69,78]. Male reproductive genes in *Drosophila*, particularly SFPs, are a classic example of this phenomenon. However little is known about the evolutionary dynamics of female reproductive tract genes because they are not well-characterized across a broad range of species. Studies in other *Drosophila* species show that some fRT genes, particularly those that code for proteases, show elevated rates of amino acid substitution [26,79–81]. Thus, these rapidly evolving fRT genes might directly interact with male seminal proteins. However we do not observe an appreciable signature of rapid divergence in fRT genes between *D. americana* and *D. novamexicana*: most fRT genes have *ω* values that are below the genome average and below the 0.5 threshold used by studies in *D. melanogaster*. These observations can be due to differences in divergence patterns between these disparate species, but can also be due to methodological differences in identifying fRT genes; we used a strict expression bias criterion to identify fRT genes, whereas other studies in *Drosophila* used expressed sequence tags (ESTs) from fRT tissues. In addition, we found that categorizing fRT genes based on functional classes or secretion signal does not affect the overall divergence estimates reported here. Thus, our results suggest that fRT-biased genes are slow evolving relative to accessory gland-biased genes in this group and likely have distinct evolutionary divergence patterns compared to species in the *Sophophora* subgenus.

Our analysis of the post-mating transcript abundance changes shows that some of the same functional categories of genes that are up-regulated in other insect taxa are also up-regulated in *D. novamexicana* females after mating [29–34,37,42,82]. A commonly enriched functional class is proteolytic enzymes, which are thought to process peptides and activate enzymatic reactions within the female reproductive tract [70]. Another class of genes that is often up-regulated after mating is immune response genes [29,30,32,37,40,42]. Our results show that post-mating induction of immune effector genes in the female reproductive tract is largely determined by male genotype: heterospecific males induce a heightened immune response in the female reproductive tract that peaks at 6 hpm, whereas conspecific males induce a mild immune response. The downstream activation of effector immune genes such as anti-microbial peptides (AMPs) is triggered by the transcription factor, Relish, which is an NF-*κ*B Imd pathway activator of immune defenses, typically against gram-positive bacteria and fungi [74]. Relish mRNA levels significantly increase at 3 hpm after heterospecific, but not conspecific mating, suggesting that AMP activation is triggered through the Imd pathway.

The initiation of immune defenses after mating in insects has been widely regarded as a mechanism to thwart potential infection during copulation, and can potentially trigger an investment trade-off between immune defense and reproduction [83–86]. However recent evidence suggests that the magnitude and direction of post-mating immune responses varies across study systems and may have additional explanations [39,41,87–90]. In particular, several studies suggest that male ejaculate components can be “immunogenic” such that they are directly triggering immune responses in females [89]. Our results support this hypothesis, and show that post-mating immune responses are elevated as a consequence of heterospecific mating, where male seminal proteins have diverged from their conspecific counterparts. Furthermore, our results indicate that additional stress responses through alternative pathways (*e.g.* JAK/STAT) are exacerbated in the heterospecific cross and may be a consequence of divergent ejaculate components. Overall the *D. novamexicana* system provides a unique opportunity to investigate the consequences of post-mating immune activation and can provide a tractable experimental system to uncouple hypotheses of immune activation as a reproductive process or pathogen defense mechanism [87,90].

During copulation, males are thought to transfer a cocktail of sperm and SFPs, but recent work has shown that males can also transfer mRNAs that can be detected in the female’s post-mating transcriptome [35,91]. In both of the reported cases in insects, the mRNAs appear to originate from the male accessory glands as the mRNAs are derived from known SFP genes or show strong expression bias in the accessory glands. We examined transcripts that were identified as up-regulated after mating in *novamexicana* and confirmed that these were transferred during copulation. Surprisingly, we found that these appear to originate from the testes. Previous work in mammals has identified spermatozoal associated mRNAs [92,93], and these mRNAs share functional properties with mRNAs that were identified in *D. melanogaster* sperm, such as ribosomal proteins [94]. Our results reveal a distinct class of ejaculate mRNAs from testes, such as heat shock proteins, and suggest that non-sperm components of the ejaculate can originate in the testes. It is not clear if the transfer of mRNAs in the male ejaculate has functional significance, and more work is needed to rule out that this mRNA transfer simply reflects the presence of cellular debris from the testes or accessory glands.

Overall this study provides new insights into the reproductive functions of the fRT in an emerging genetic model clade, and lays the groundwork for future investigation into the genet basis of gamete interactions in *Drosophila*

## Acknowledgments

We thank Amanda Manfredo for technical assistance, Sofie Delbare for discussion and comments on the manuscript, the Cornell Biotechnology Resource Center for sequencing and computational resources, and members of the Clark and Wolfner labs for comments and suggestions. This work was supported by NIH grant R01-HD059060 to A.G.C and M.F.W.

## Supporting Information

**Figure S1.**
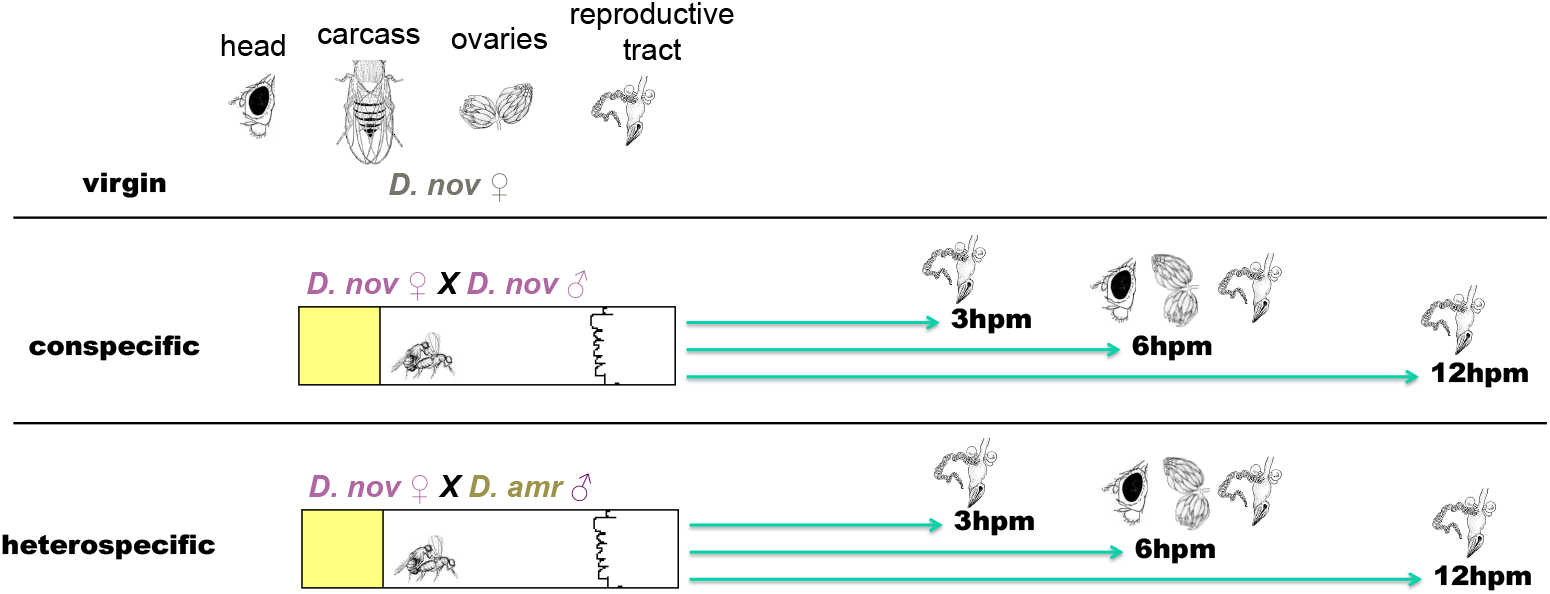
Experimental design and tissues used in the study. Three tissues were used to generate short read RNA-seq data: heads, ovaries, and lower reproductive tracts. We also used the gonadectomized carcass to identify tissue-biased transcripts. Virgin RNA-seq libraries included all four tissue samples, while post-mating libraries included heads (6 hpm), ovaries (6 hpm), and lower reproductive tracts (3 hpm, 6 hpm, and 12 hpm).

**Figure S2.**
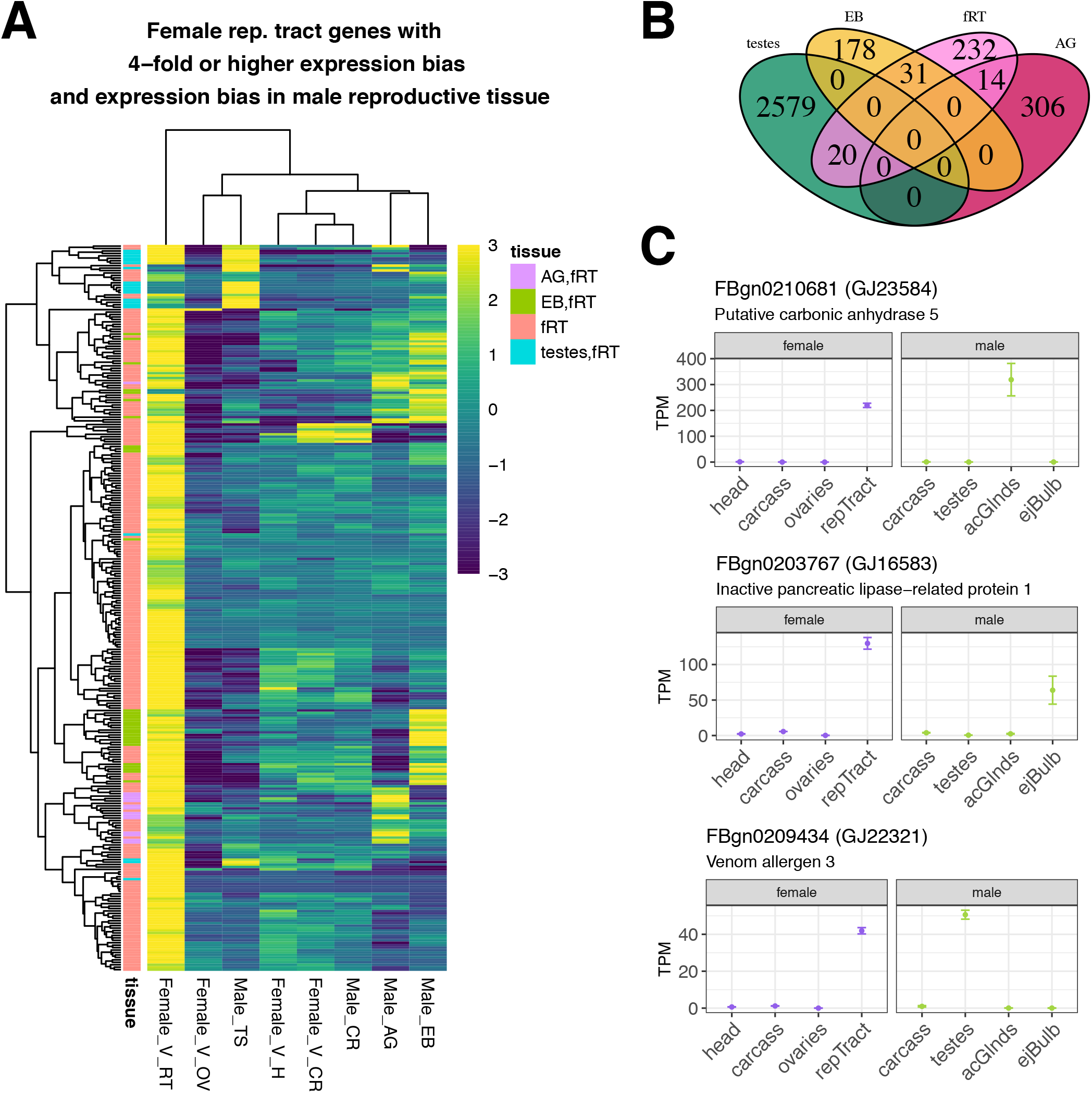
Shared reproductive tissue-biased genes in females and males. **(A)** Heatmap of centered log2 mean TPM of fRT genes (logFC ≥2) across male and female *D. novamexicana* tissues. Row annotations on the left indicate the tissue-biased classification with the key on the right (fRT: female reproductive tract; AG: accessory glands; EB: ejaculatory bulb). **(B)** Venn diagram showing the overlap of fRT-biased genes with male reproductive tract-biased genes. **(C)** Examples of three genes that show fRT-biased expression in females but also show accessory gland-biased (top), ejaculatory bulb-biased (middle), or testes-biased (bottom) expression in males.

**Figure S3.**
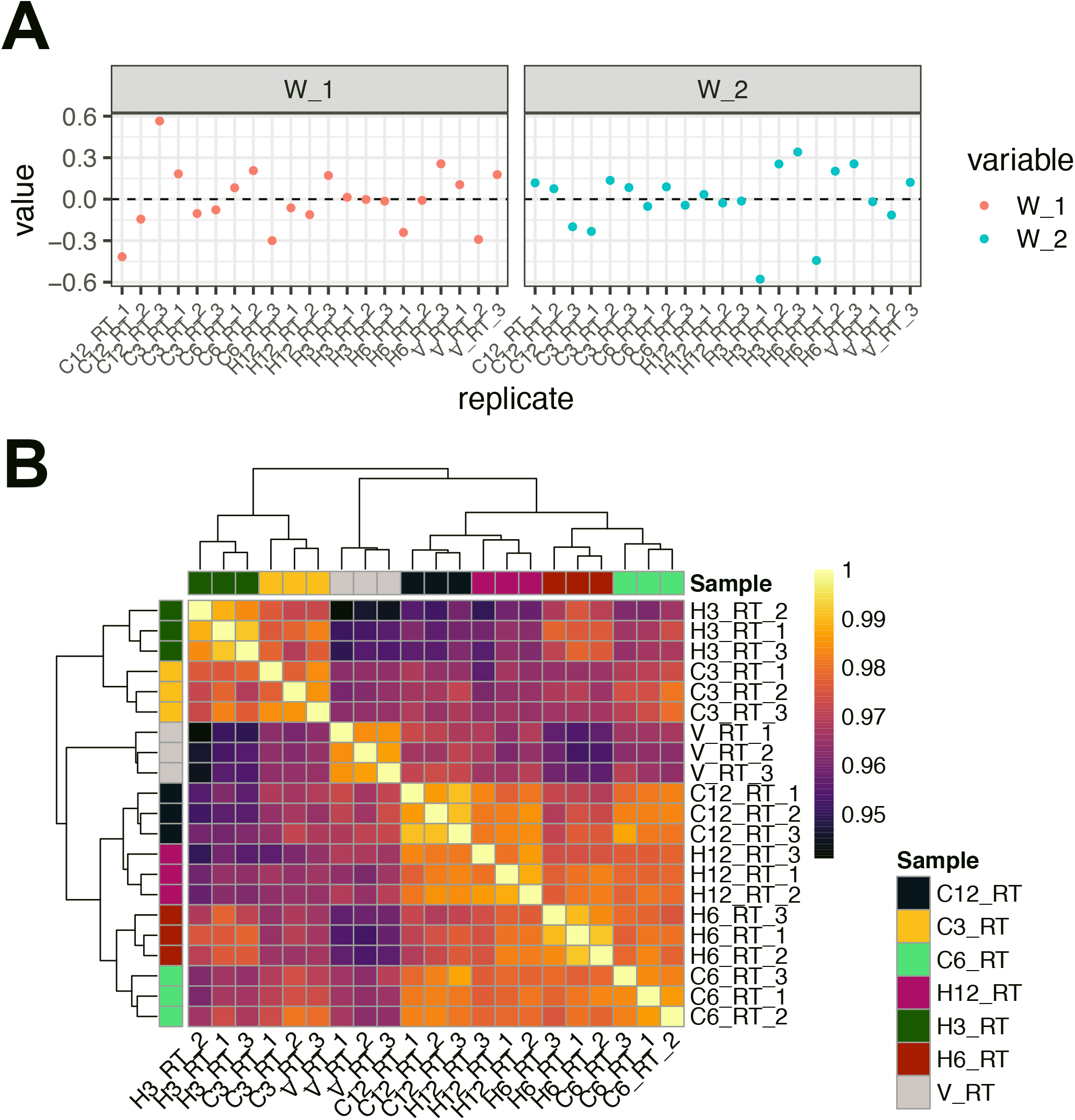
**(A)** Distribution of RUVseq residual correction variables across fRT replicates and samples with *k* = *2*. **(B)** Correlation matrix (Pearson coefficient) of virgin and post-mating fRT samples. Row and column annotation bars represent sample ID.

**Figure S4.**
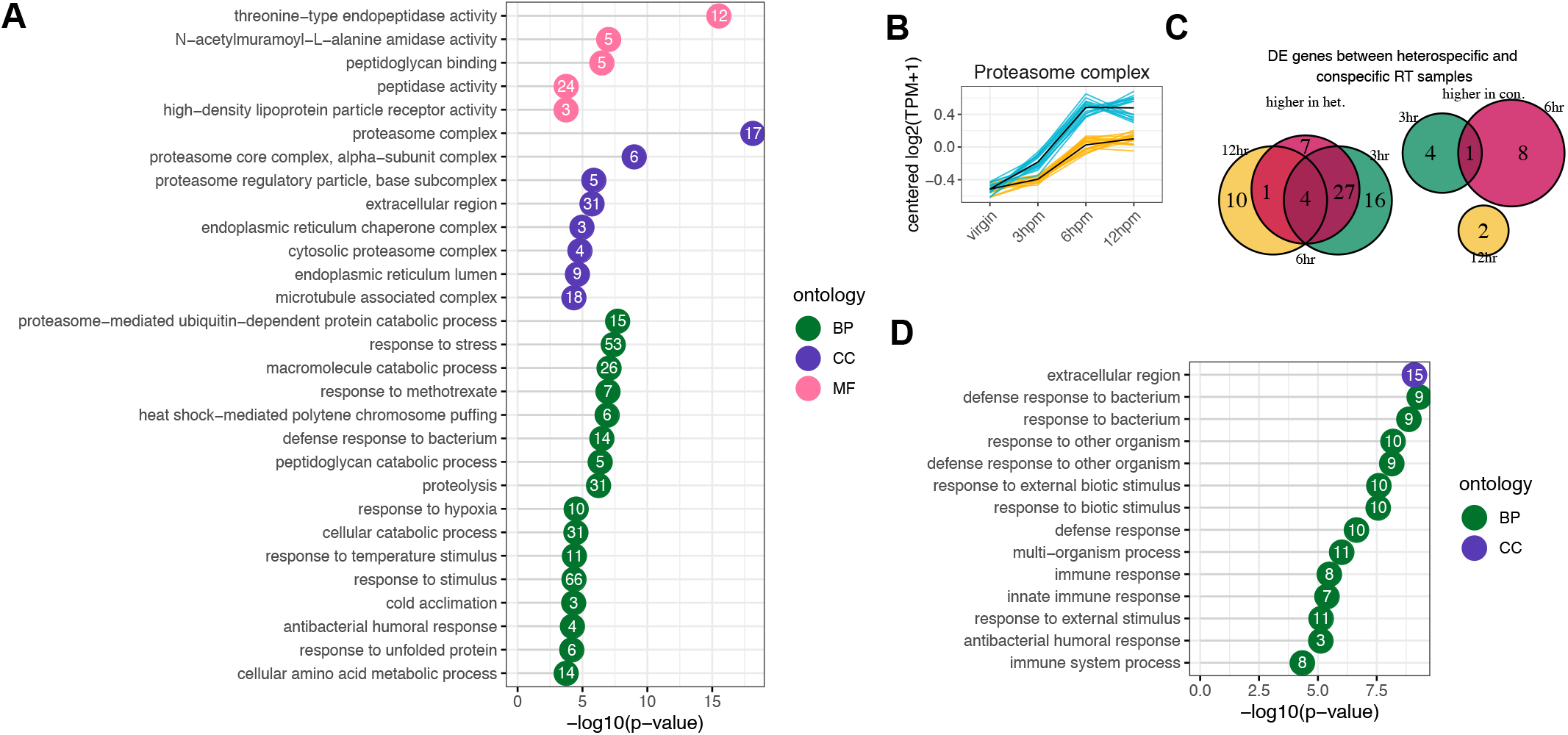
**(A)** Significantly enriched Gene Ontology (GO) terms among up-regulated genes in the fRT after mating. Redundant and/or nested GO terms were trimmed using GOtrim. The number of differentially abundant transcripts that belong to each ontology term is indicated within the circle, and the ontology category is indicated by color (BP: Biological Process; CC: Cellular Component; MF: Molecular Function). **(B)** Genes that code for components of the core proteasome complex that are up-regulated in the female fRT after mating to conspecific (yellow) or heterospecific (blue) males. **(C)** Venn diagram of the number of transcripts that are differentially abundant between conspecific and heterospecific fRT post-mating samples. **(D)** Significantly enriched GO terms among genes that have significantly higher abundance in the heterospecific fRT samples compared to the conspecific fRT samples (GO terms not pruned as in (A)).

**Figure S5.**
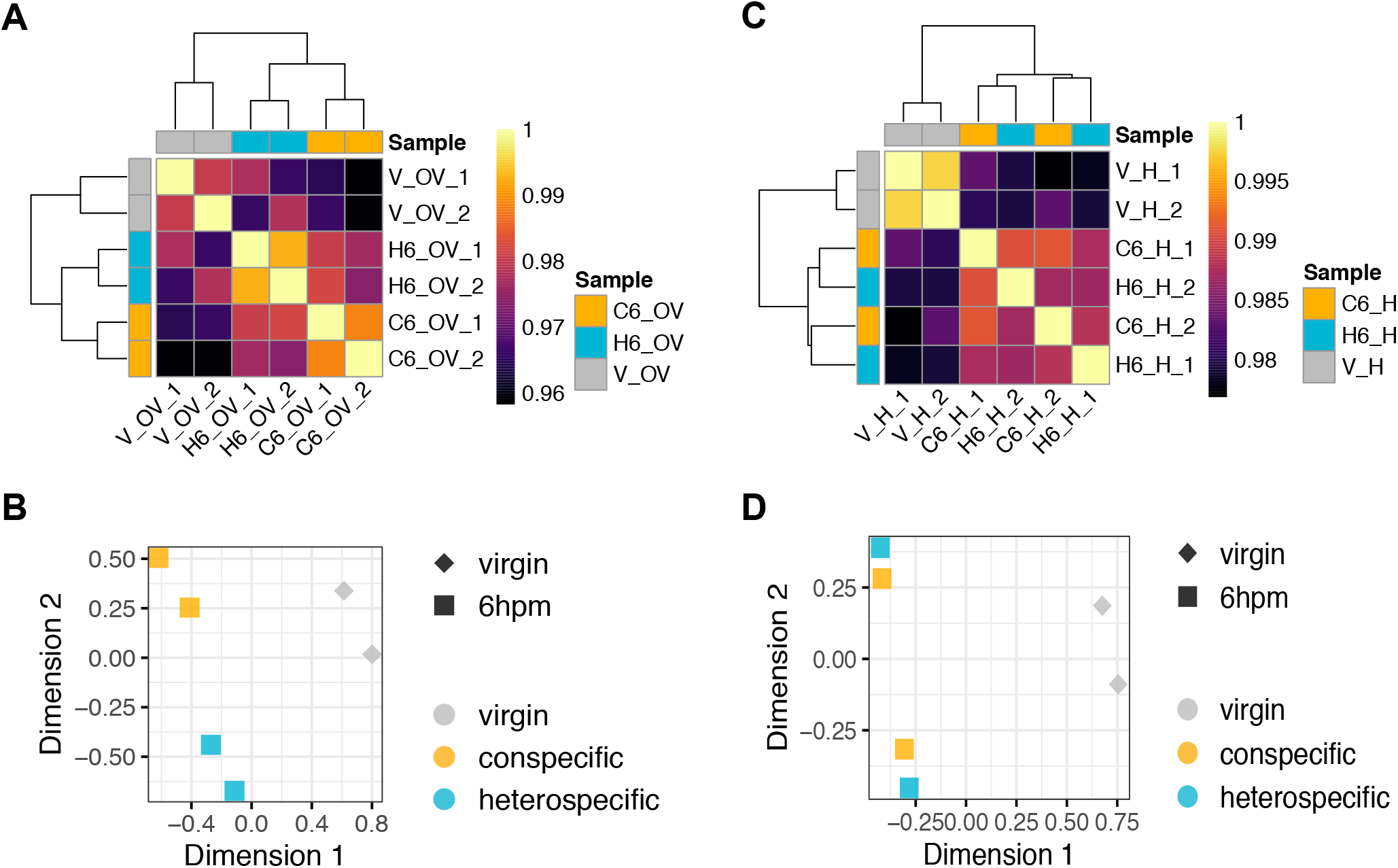
**(A)** Correlation matrix (Pearson coefficient) of virgin and post-mating ovary samples. **(B)** MDS plot of virgin and post-mating ovary samples. **(C)** Correlation matrix (Pearson coefficient) of virgin and post-mating head samples. **(D)** MDS plot of virgin and post-mating head samples.

